# Transcription factors GAF and HSF act at distinct regulatory steps to modulate stress-induced gene activation

**DOI:** 10.1101/055921

**Authors:** Fabiana M. Duarte, Nicholas J. Fuda, Dig B. Mahat, Leighton J. Core, Michael J. Guertin, John T. Lis

## Abstract

The coordinated regulation of gene expression at the transcriptional level is fundamental to organismal development and homeostasis. Inducible systems are invaluable when studying transcription because the regulatory process can be triggered instantaneously, allowing the tracking of ordered mechanistic events. Here, we use Precision Run-On sequencing (PRO-seq) to examine the genome-wide Heat Shock (HS) response in *Drosophila* and the function of two key transcription factors on the immediate transcription activation or repression of all genes regulated by HS. We identify the primary HS response genes and the rate-limiting steps in the transcription cycle that GAGA-Associated Factor (GAF) and HS Factor (HSF) regulate. We demonstrate that GAF acts upstream of promoter-proximally paused RNA Polymerase II (Pol II) formation, likely at the step of chromatin opening, and that GAF-facilitated Pol II pausing is critical for HS activation. In contrast, HSF is dispensable for establishing or maintaining Pol II pausing, but is critical for the release of paused Pol II into the gene body at a subset of highly-activated genes. Additionally, HSF has no detectable role in the rapid HS-repression of thousands of genes.

## Introduction

The <u>H</u>eat <u>S</u>hock (HS) response in *Drosophila melanogaster* has been an effective model system to discover and study mechanisms of transcription and its regulation (Guertin et al. 2010). This highly conserved protective mechanism (Lindquist and Craig 1988) is regulated at the transcriptional level by the <u>H</u>eat <u>S</u>hock transcription <u>F</u>actor (HSF) (Wu 1995). When activated by stress, HSF potently activates expression of HS genes, resulting in the accumulation of molecular chaperones, the <u>H</u>eat <u>S</u>hock <u>P</u>roteins (HSPs), which helps the cell cope with stress-induced protein aggregation and misfolding (Lindquist and Craig 1988; Schlesinger 1990).

The transcriptional HS response has been studied largely using *Hsp70* as a model gene (Guertin et al. 2010). *Hsp70* maintains a promoter-proximally paused RNA <u>Pol</u>ymerase <u>II</u> (Pol II) molecule 20-40 bp downstream of the *T*ranscription <u>S</u>tart <u>S</u>ite (TSS) that is released to transcribe the gene at a low level during normal non-stress conditions (Rougvie and Lis 1988; Rasmussen and Lis 1993). The transcription factor <u>G</u>AGA <u>A</u>ssociated <u>F</u>actor (GAF) is bound to the promoter of *Hsp70* prior to HS, and GAF is important for the establishment and stability of paused Pol II (Lee et al. 1992; Wang et al. 2005; Lee et al. 2008; Kwak et al. 2013). GAF has a key role in keeping the promoter region open and free of nucleosomes (Tsukiyama et al. 1994; Okada and Hirose 1998; Fuda et al. 2015), which allows the recruitment of general transcription factors and the initiation of transcription by Pol II. Upon HS induction, HSF trimerizes and is rapidly recruited to the promoter, where it binds to its cognate <u>HS</u> DNA <u>E</u>lements (HSEs) (Pelham 1982; Xiao and Lis 1988). After binding, HSF directly and indirectly recruits coactivators and other factors (Lis et al. 2000; Saunders et al. 2003; Smith et al. 2004; Ardehali et al. 2009) that affect the chromatin structure and composition, and promotes the release of Pol II from the paused complex into productive elongation. This transition from the paused state into productive elongation depends critically on the positive elongation factor P-TEFb, and has been shown to be a very general step that is essential for the regulation of virtually all genes across different species (Rahl et al. 2010; Jonkers et al. 2014). The net result of this molecular cascade is an increase in transcription levels that can be ~200-fold for some of the HS-regulated genes (Lis et al. 1981).

Although the independent mechanisms of promoter-proximal pausing and escape to productive elongation have been well studied in the context of HS activation of *Hsp70*, we lack a comprehensive characterization of the genome-wide changes in transcription that result from HS. A thorough characterization of the affected genes is necessary to determine the generality and diversity of the roles of transcription factors such as GAF and HSF in the HS response and provide the statistical power to assess mechanisms of transcription regulation.

Previous studies have mapped HSF binding sites during normal growth conditions and after HS, and observed that HSF recruitment to a promoter is neither necessary nor sufficient to direct HS gene activation (Trinklein et al. 2004; Guertin and Lis 2010; Gonsalves et al. 2011). Nonetheless, the rules governing the specificity of activation and repression across the *Drosophila* genome remain incomplete. Transcriptional changes after HS have also been measured in *Drosophila* and other organisms (Leemans et al. 2000; Guhathakurta et al. 2002; Murray et al. 2004; Trinklein et al. 2004; Sørensen et al. 2005; Gonsalves et al. 2011; Vihervaara et al. 2013); however, these studies were limited in resolution both temporally and spatially by measuring steady-state levels of mature mRNA. Furthermore, measurement of mRNAs cannot distinguish effects on mRNA stability (Lindquist and Petersen 1990) and pre-mRNA processing (Yost and Lindquist 1986; Shalgi et al. 2014) from transcription, or primary from secondary effects of the HS response.

To overcome these limitations, we queried the genome-wide distribution of transcriptionally-engaged RNA polymerases before and after HS induction using the <u>P</u>recision nuclear <u>R</u>un-<u>O</u>n and sequencing (PRO-seq) assay and quantified differentially expressed genes. PRO-seq has high sensitivity and high spatial and temporal resolution, providing an unprecedented comprehensive view of the transcriptional profiles of cell populations. We show that the HS response is rapid and pervasive, with thousands of genes being repressed after 20 minutes of HS and hundreds of genes being activated; moreover, the activated genes are not limited to the classical HSP genes. Promoter-proximal pausing is highly prevalent among the activated genes prior to HS, and here we demonstrate that its establishment on a subset of genes is dependent on GAF binding upstream and proximal to the TSS. Moreover, GAF depletion abrogates pausing and consequently impairs HS activation, indicating that this step in early transcription elongation is essential for gene activation. We also identify two factors that correlate with Pol II pausing and HS activation at genes that exhibit GAF-independent pausing. Furthermore, we demonstrate that only a relatively small fraction of HS activated genes are regulated by HSF, and that depletion of HSF does not affect pausing levels before or after HS. This study provides a genome-wide view of HS-induced transcriptional regulation and an understanding of how promoter context affects this process.

## Results

### *Drosophila* transcriptional Heat Shock response is rapid and pervasive

We measured nascent transcription levels by PRO-seq in *Drosophila* S2 cells prior to HS (<u>N</u>on-<u>H</u>eat <u>S</u>hock – NHS) and 20 minutes after an instantaneous and continuous HS stress (Figure 1A, Table S1). PRO-seq maps the active sites of transcriptionally engaged RNA polymerase complexes by affinity-purification and sequencing of nascent RNAs after a terminating biotin-NTP is incorporated during a nuclear run-on experiment (Kwak et al. 2013). The density of sequencing reads is proportional to the number of transcriptionally-engaged polymerase molecules present at each position when the nuclei were isolated. PRO-seq has base pair resolution, is strand specific, and is not affected by the background levels of accumulated RNAs (Kwak et al. 2013). Biological replicates were highly correlated for both promoter and gene body PRO-seq reads (Spearman's coefficient ranged between 0.96-0.99, Figure S1A-B, left panels). The expected genome-wide changes in transcription that occur during HS made it unfeasible to use total number of reads to normalize our datasets between conditions. Therefore, we normalized our libraries using a set of genes previously shown to have the same RNA Pol II ChIP-seq signals in NHS and HS *Drosophila* S2 cells where consistent backgrounds of ChIP-seq provide a basis of normalization (Teves and Henikoff 2011) (see Materials and Methods for the normalization method and our validation tests).

**Figure 1:**
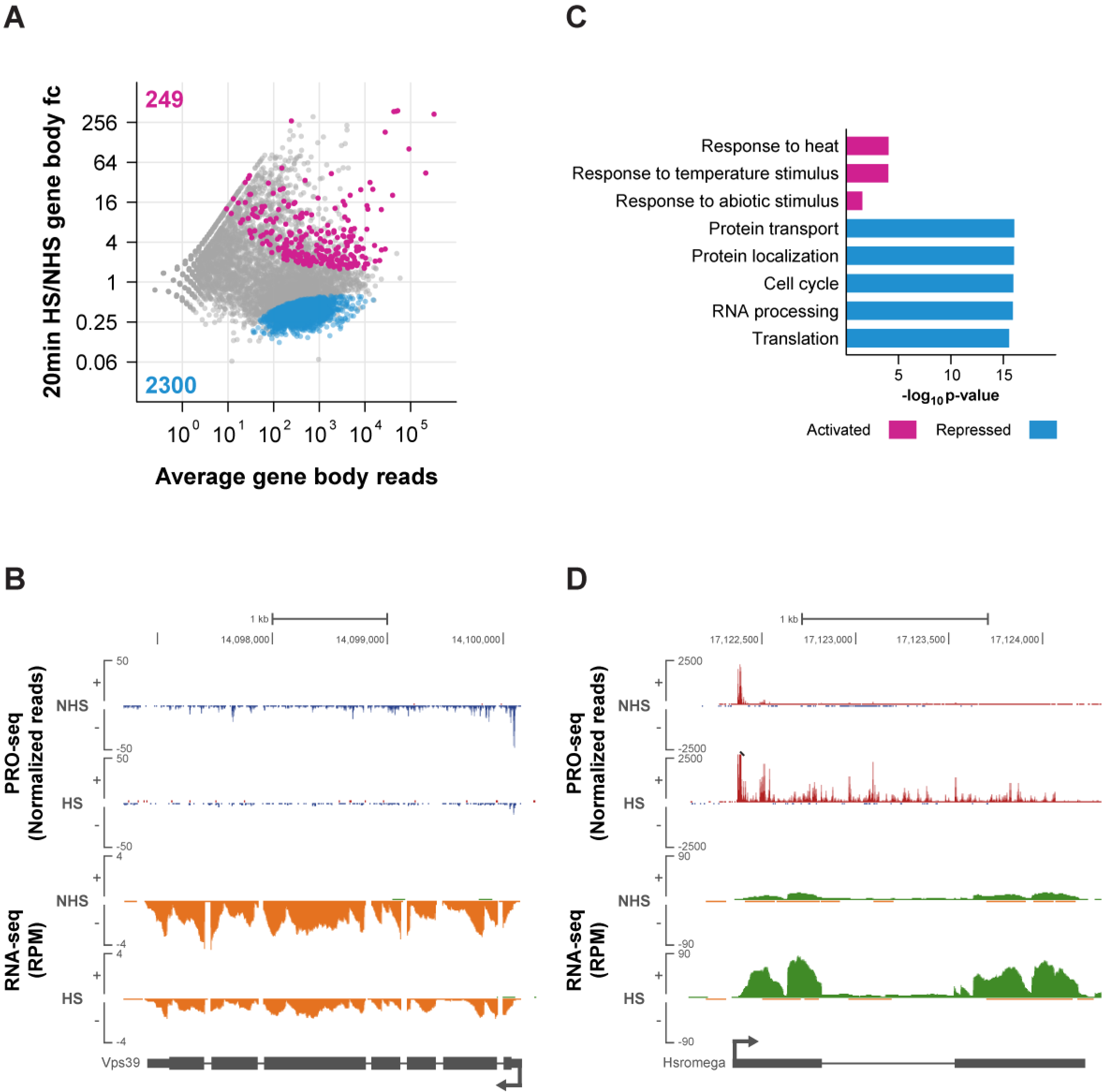
*Drosophila* transcriptional <u>H</u>eat <u>S</u>hock response is rapid and pervasive. **(A)** DESeq2 analysis of PRO-seq gene body reads between 20min HS-treated and NHS cells displayed as an MA plot. Significantly changed genes were defined using an FDR of 0.001. Activated genes are labeled in magenta and repressed genes in blue. The number of genes in each class is shown in the plot. fc = fold-change. **(B)** Representative view of a HS repressed gene in the UCSC genome browser (Kent et al. 2002). PRO-seq normalized reads for the plus strand are shown in red and for the minus strand in blue. RNA-seq reads for the plus strand are shown in green and for the minus strand in orange. Gene annotations are shown at the bottom. **(C)** Gene ontology terms enriched in the HS activated and repressed classes. **(D)** Representative view of a HS activated gene in the UCSC genome browser (Kent et al. 2002). Axes are the same as in B.

We used DESeq2 (Love et al. 2014) to identify genes whose gene body reads significantly change after HS, using an FDR of 0.001 (Table S2). We observed a widespread shutdown of transcription, with 2300 genes being significantly repressed after HS (Figure 1A, blue points; Figure 1B has an example of a repressed gene – *Vps39*). This finding is in agreement with low resolution studies in *Drosophila* salivary gland polytene chromosomes that have shown that total Pol II levels and transcription decrease in response to HS (Spradling et al. 1975; Jamrich et al. 1977). A previous Pol II ChIP-seq study in *Drosophila* S2 cells has also observed a genome-wide decrease of Pol II levels in gene bodies (Teves and Henikoff 2011). Not surprisingly, measurements of steady-state mRNA levels before and after HS, including micro-array studies and our own RNA-seq data (Figure S2, Table S3), were unable to detect a genome-wide shutdown of transcription, despite having the sensitivity to detect a decrease in mRNA levels for some genes (Figure 1B). Measurements of mRNA do not detect genome-wide transcriptional repression because the reduction of mRNA levels are obscured by steady-state levels of mRNAs already present in the cells; these mRNAs have much longer half-lives than the short HS time points examined herein. Overall, our results greatly expand upon these previous studies, identifying and quantifying the individual genes whose transcription is repressed after HS using a base-pair resolution method that specifically maps transcriptionally-engaged RNA polymerase molecules.

**Figure 2:**
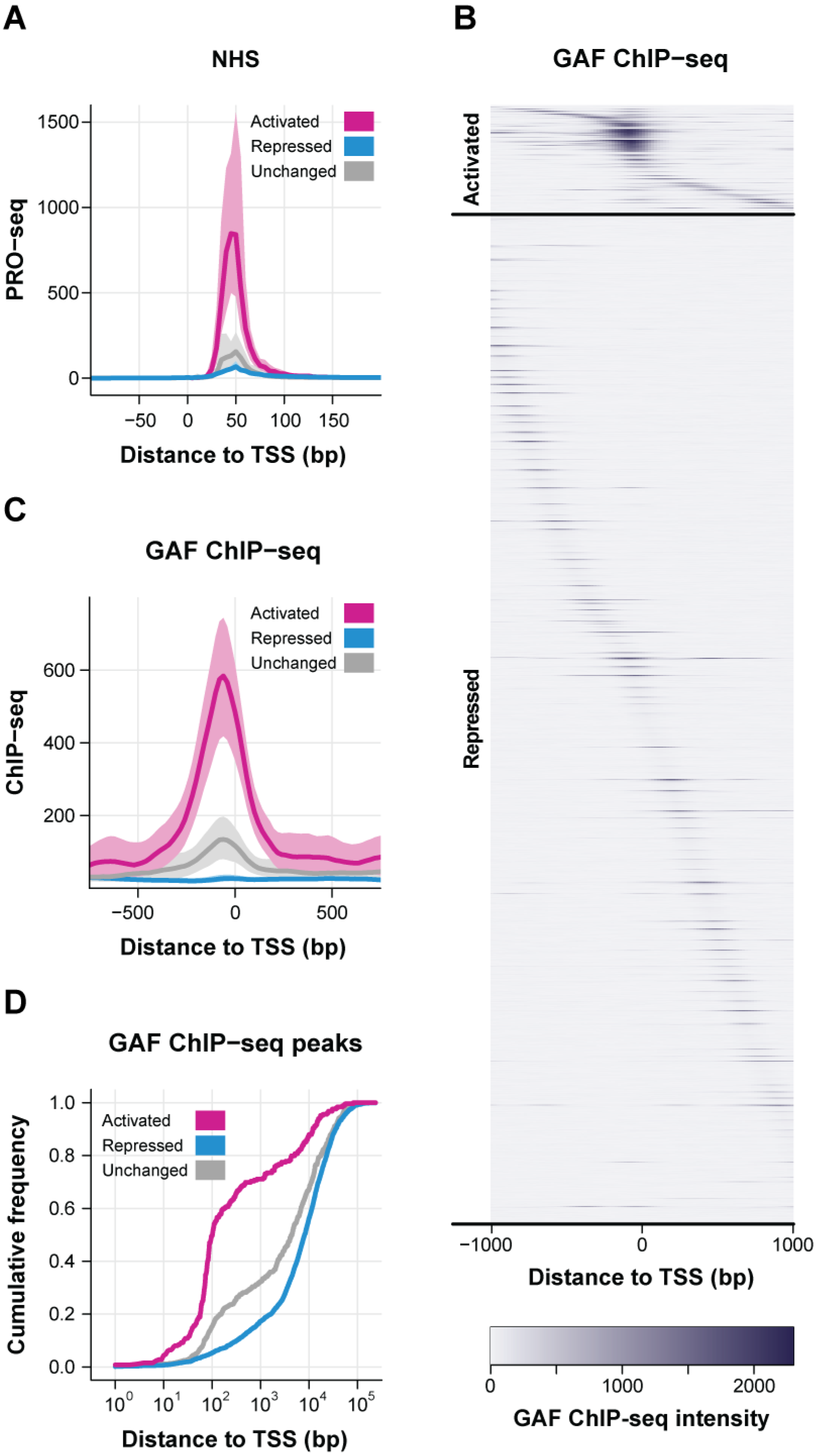
GAGA factor is highly enriched in the promoter region of HS activated genes prior to HS. **(A)** Average normalized PRO-seq reads between −100 to +200 bp to the TSS (in 5 bp bins) for the LacZ-RNAi NHS dataset of HS activated (n=249), repressed (n=2300), and unchanged (n=517) genes. The shaded area represents the 75% confidence interval. **(B)** Heatmap showing the GAF ChIP-seq signal in 20bp windows from ±1 kb to the TSS of HS activated (n=249) and repressed (n=2300) genes. For each class, genes were ordered by the distance between the highest intensity window and the TSS. **(C)** Average GAF ChIP-seq reads between −750 to +750 bp to the TSS (in 20 bp bins) of HS activated (n=249), repressed (n=2300), and unchanged (n=517) genes. The shaded area represents the 75% confidence interval. **(D)** Cumulative distribution plots of the distance between the closest GAF ChIP-seq peak and the TSS of each gene in the HS activated, repressed, and unchanged classes.

Gene ontology (GO) analysis reveal that the HS repressed class is enriched for genes involved in basic cellular processes, such as cell cycle, RNA processing, protein transport and localization and translation (Figure 1C). This is consistent with previous findings in mammalian cells (Murray et al. 2004; Trinklein et al. 2004), and is expected as cells enter into a defensive non-growth condition triggered by HS stress.

Although not as abundant as the repressed class, hundreds of genes are activated by HS, many very highly (Figure 1A, magenta points; Figure 1D has an example of an activated gene – *Hsromega*). Notably, we find that all 7 classical HSP genes show strong inductions after 20 minutes HS and are among the top 10 genes with highest HS induction (Table S2), with fold-changes ranging from 44 to 384-fold. Consistent with this result, GO analysis reveal that the HS activated class is enriched for genes involved in the response to temperature and abiotic stimuli (Figure 1C). Besides the classical HSP genes, our data reveal the activation of many genes that were not previously associated with the HS response and provide a comprehensive characterization and quantification of genes whose transcription is directly activated by HS.

We measured nascent transcription levels as a function of time after HS induction to determine how fast activated and repressed genes respond to HS (Figure S3, Table S1). Biological replicates produced high correlations for PRO-seq reads within either the promoter or gene body regions (Spearman's coefficient ranged between 0.9-0.99, Figure S1C-D). The sequential HS time points displayed a progressive increase in the number of genes that were significantly activated (Figure S3A, magenta points; S3B has an example of an activated gene – *CG13321*) and repressed (Figure S3A, blue points; S3C has an example of a repressed gene – *CG14005*) by HS. No genes are significantly different after 30 seconds and only a small number of genes are significantly different after 2 minutes of HS. We observe a substantial genome-wide response to HS as early as 5 minutes post-HS, the response is even more pervasive at later time points. Previous studies have shown that classical HSP genes are activated very rapidly (O'Brien and Lis 1993; Zobeck et al. 2010), but herein we demonstrate that many other genes have a rapid response, both for activation and repression. The number of significantly activated and repressed genes further increases after 10 and 20 minutes of HS (Figure S3A). Overall, our results demonstrate that the HS response produces an immediate and primary change in the transcription levels of 30% of the unambiguously mappable mRNA encoding genes (Core et al. 2012), with the repression of thousands of genes and the activation of many hundreds of genes.

### Activated genes are highly paused prior to HS

During normal cell growth, classical HSP genes have a paused, transcriptionally engaged polymerase between 20-50 bp downstream of the TSS (Rougvie and Lis 1988; Rasmussen and Lis 1993). Furthermore, promoter-proximal Pol II pausing is the major regulatory step for the HS activation of the *Hsp70* gene, where it maintains the promoter region open and accessible to transcription factors (Lee et al. 1992; Shopland et al. 1995). We used our PRO-seq data to determine if promoter-proximal pausing is a common feature of HS activated genes. The average PRO-seq read intensity profile across HS activated genes reveals a strong peak in the promoter-proximal region, which is substantially higher than repressed or unchanged gene classes (Figure 2A). DNase I hypersensitivity data (Kharchenko et al. 2011) indicates that the promoter region of HS activated genes is more accessible than the other two classes under basal uninduced conditions (Figure S4A). This data is consistent with the notion that promoter-proximal pausing is important to maintain an open chromatin environment around the TSS.

We calculated the pausing index (PI), which is the ratio of read density in the promoter-proximal region relative to the gene body, for each individual gene (Core et al. 2008) (Table S4). The vast majority of HS activated genes (~90%) were classified as paused (Fisher's exact p-value <= 0.01, Figure S4B) (Core et al. 2008). The PI is significantly higher for activated genes compared to the repressed (Mann-Whitney *U* test p-value < 2.2 × 10^-16^) and unchanged classes (Mann-Whitney *U* test p-value < 2.2 × 10^−16^) (Figure S4C). Although the pausing levels of HS repressed genes are not as high as the activated class (Figure 2A), a considerable percentage of repressed genes were also classified as paused (~80%, Figure S4B). Overall, our results indicate that high levels of promoter-proximal pausing are a general feature of HS-induced genes prior to HS and may play an important role in poising these genes for HS activation by transcription factors.

### GAGA factor is highly enriched in the promoter region of HS activated genes

To identify candidate factors that play a role in allowing genes to be HS-activated, we screened modENCODE and other publically available genomic transcription factor binding data for factors that are differentially enriched in HS activated relative to repressed or unchanged genes prior to HS. The most striking differential enrichment was observed for GAF (Figure 2B, GAF ChIP-seq data from Fuda et al. 2015). As seen in Figure 2B, when compared to the repressed class, HS activated genes show enriched GAF binding immediately upstream of the TSS, which is also evidenced by a peak in the average ChIP-seq intensity profile (Figure 2C). Furthermore, *de novo* motif analysis identified the DNA sequence bound by GAF, the GAGA element (Omichinski et al. 1997; Wilkins 1998), as the most significantly overrepresented motif in the region immediately upstream of the TSS of HS activated genes (Figure S4D).

We then identified the closest GAF ChIP-seq peak to the TSS of each gene and plotted the cumulative distribution of these distances for our three gene classes (activated, repressed and unchanged) (Figure 2D). GAF binds significantly closer to activated genes than the repressed (Figure 2D, Kolmogorov-Smirnov test p-value < 2.2 × 10^-16^) and unchanged classes (Figure 2D, Kolmogorov-Smirnov test p-value < 2.2 × 10^-16^), and over 70% of activated genes are bound by GAF within ±1kb of the TSS. These results suggest that GAF binding close to the TSS prior to HS is important for the activation of HS-induced genes.

### GAF is critical for HS activation when bound immediately upstream of the core promoter

To investigate whether GAF binding is essential for HS activation, we performed PRO-seq in biological replicates of GAF-RNAi treated cells prior to HS and after 20 minutes of HS (Figure 3A–B, Table S1) (Spearman's coefficient ranged between 0.96-0.99, Figure S1A-B, right panels). The decrease in GAF protein levels after the RNAi treatment produced similar numbers of genes that were significantly activated or repressed by HS (Figure S5A, compare to Figure 1A); however, comparison of the HS gene body reads in the GAF-RNAi and LacZ-RNAi control identified many genes that were significantly affected by the knockdown. The HS PRO-seq levels of 20% of activated genes were affected by GAF-RNAi and nearly all were reduced (Figure 3C, left panel), while less than 1% of the repressed class were affected (Figure 3C, right panel), demonstrating that GAF is important for HS activation, but not for repression. Greater than 90% of the genes that have reduced HS induction after GAF knockdown have GAF binding within ±1kb of the TSS (Figure 3C, left panel). Taken together, these results indicate that promoter-bound GAF is indispensable for the proper activation of many HS activated genes.

**Figure 3:**
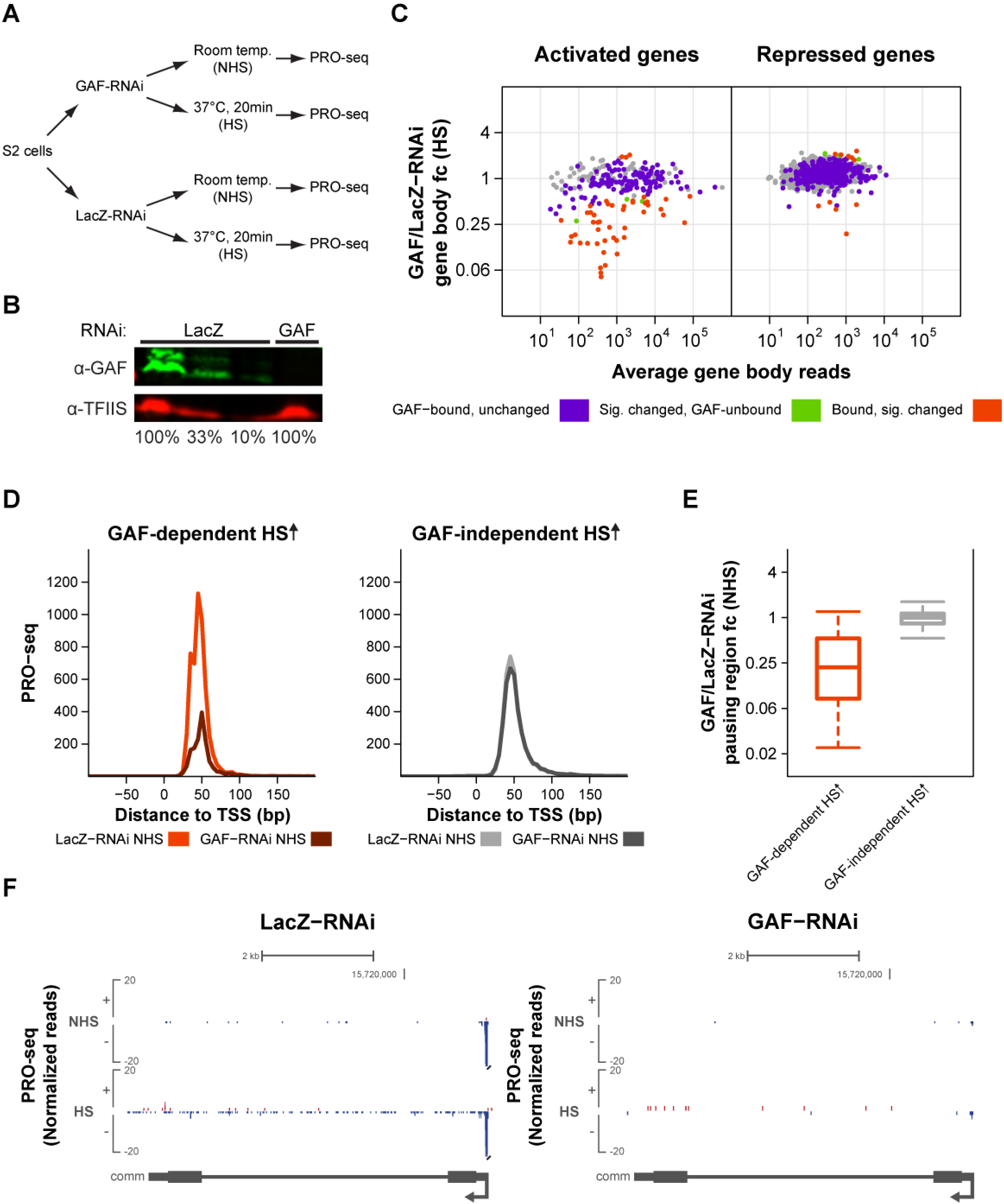
GAF's role in HS activation correlates with its function in establishing promoter-proximal pausing prior to HS. **(A)** Experimental set-up. *Drosophila* S2 cells were treated with either GAF or LacZ RNAi for 5 days. Nuclei were then isolated for PRO-seq after cells were incubated at room temperature (NHS) or <u>H</u>eat <u>S</u>hocked for 20 minutes (HS). **(B)** Western blot of whole cell extracts from LacZ and GAF-RNAi treated cells for GAF and TFIIS (loading control). 100% is equivalent to 1.5 × 10^6^ cells. **(C)** DESeq2 analysis to determine the effect of GAF-RNAi treatment on the PRO-seq gene body reads after HS for the HS activated (n=249) and repressed (n=2300) classes. DESeq2 was used to identify significantly changed genes between GAF-RNAi HS and LacZ-RNAi HS cells and the results are displayed as MA plots. Significantly changed genes were defined using an FDR of 0.001. GAF-bound genes are labeled in purple, significantly changed genes (according to DESeq2) are labeled in green and genes that are both GAF-bound and significantly changed are labeled in orange. fc = fold-change. **(D)** Average normalized PRO-seq reads between −100 to +200bp to the TSS (in 5 bp bins) for the LacZ-RNAi NHS and GAF-RNAi NHS treatments of genes with GAF-dependent (n=44) or GAF-independent (n=199) HS activation. **(E)** Box-plot showing the GAF/LacZ-RNAi pausing region fold-change prior to HS (NHS) for genes with GAF-dependent or GAF-independent HS activation. Over 70% of the genes with GAF-dependent HS activation have significantly reduced pausing upon GAF depletion prior to HS, while only 15% of the GAF-independent genes were significantly affected. **(F)** Representative view in the UCSC genome browser (Kent et al. 2002) of a gene with GAF-dependent pausing prior to HS whose activation is inhibited by GAF-RNAi treatment. PRO-seq normalized reads for the different RNAi treatments (LacZ and GAF) before and after HS for the plus strand are shown in red and for the minus strand in blue. Gene annotations are shown at the bottom.

GAF is critical for the HS activation of many genes; however, the induction of over 70% of the genes that are bound by GAF prior to HS is not affected by GAF knockdown. These two classes of GAF-bound genes, which respond differentially to GAF knockdown, cannot simply be explained by differences in the response to the RNAi treatment, since the GAF ChIP-seq signal for both classes is similarly reduced by the knockdown (Figure S5B). However, GAF-bound genes with GAF-dependent HS activation have significantly higher GAF ChIP-seq intensities when compared to the GAF-bound genes with GAF-independent HS activation (Figure S5C, Mann-Whitney *U* test p-value = 8.96 × 10^−10^). The class of GAF-bound, HS-activated genes whose induction is dependent on GAF has a strong preference for GAF binding immediately upstream of the TSS, between −100 to −50 bp (Figure S5D, right panel). Taken together, these results suggest that higher binding levels and positioning upstream and proximal to the TSS are essential for GAF's role in HS activation.

### GAF's role in HS activation correlates with its function in establishing promoter-proximal pausing prior to HS

GAF has been shown to have a role in the establishment of promoter-proximal pausing and consequent HS activation of two classical HSP genes (Glaser et al. 1990; Lee et al. 1992; Lu et al. 1993; O'Brien et al. 1995), and a recent study has demonstrated that pausing was significantly reduced on a large subset of GAF-bound genes upon GAF depletion (Fuda et al. 2015). However, the role of GAF-mediated pausing in gene activation has not yet been studied in a comprehensive genome-wide manner. We hypothesize that GAF's role in HS activation is connected to its ability to create promoter-proximal pausing prior to HS.

To test this hypothesis, we compared the NHS promoter-proximal PRO-seq reads for the LacZ-RNAi control and GAF-RNAi treatment between the subset of GAF-bound genes whose HS induction is dependent on GAF (GAF-dependent HS activation) and the HS activated genes whose induction is unaffected by GAF depletion (GAF-independent HS activation). As observed in Figure 3D, there is a substantial reduction in the NHS pausing levels after GAF knockdown for genes with GAF-dependent HS activation, while the NHS pausing levels of the GAF-independent class are largely unaffected. To quantify this effect, we compared the LacZ-RNAi and GAF-RNAi NHS reads in the pausing region for genes with GAF-dependent or GAF-independent HS activation. As observed in Figure 3E, most genes with GAF-dependent HS activation have reduced number of reads (fold-change < 1) in the pausing region upon GAF knockdown prior to HS. In contrast, the distribution of fold-changes for the GAF-independent class is centered around 1, indicating that GAF binding prior to HS is not essential to establish pausing at these genes. Figure 3F has an example of a HS-activated gene that displays GAF-dependent pausing prior to HS whose induction is inhibited by GAF knockdown. Taken together, these results indicate that GAF's role in HS activation strongly correlates with its function in establishing promoter-proximal pausing prior to HS.

### Insulator proteins and M1BP are enriched in the promoter region of HS activated genes with GAF-independent induction

While nearly all activated genes display promoter-proximal pausing prior to HS, we have shown that GAF is essential for pausing establishment and HS activation on a subset of these genes. To identify factors that can contribute to the establishment of pausing on GAF-independent genes, we screened modENCODE and other publically available chromatin factor and histone modification ChIP-seq or ChIP-chip datasets for factors that are differentially enriched in the promoter region of GAF-independent relative to GAF-dependent genes prior to HS. The most striking differential enrichment was observed for the transcription factor <u>M</u>otif <u>1</u> <u>B</u>inding <u>P</u>rotein (M1BP) (ChIP-seq data from Li and Gilmour 2013), the insulator protein BEAF-32 (ChIP-chip data from Schwartz et al. 2012), and the chromodomain containing protein Chromator (ChIP-chip data from Kharchenko et al. 2011) (Figure 4A–C). M1BP is a recently discovered zinc-finger transcription factor that has been shown to orchestrate promoter-proximal pausing in a GAF-independent manner (Li and Gilmour 2013). BEAF-32 is one of the insulator associated proteins identified in *Drosophila* (Zhao et al. 1995), and Chromator, which was initially identified as a mitotic spindle protein (Rath et al. 2004) and later implicated in the regulation of chromosome structure through partial cooperation with BEAF-32 (Rath et al. 2006; Gan et al. 2011), and both of these proteins are enriched at the boundaries of physical chromosomal domains (Hou et al. 2012; Sexton et al. 2012). Remarkably, almost no overlap exists between genes with GAF-dependent HS activation and genes bound by M1BP or insulator proteins within ±1kb of the TSS (Figure 4D). The mutually exclusive distributions of GAF and M1BP in promoter-proximal pausing has been previously reported (Li and Gilmour 2013), and our results suggest a possible role for M1BP in pausing and HS activation. Similarly to M1BP, the mutually exclusive distribution of insulator proteins and the GAF-dependent subset suggests that BEAF-32 and/or Chromator may have a role in generating promoter-proximal pausing when bound proximally to the TSS. GAF has also been classified as an insulator protein with enhancer-blocking activity (Ohtsuki and Levine 1998; Schweinsberg et al. 2004), which suggests a possible overlap between insulator function and a role in maintaining an open chromatin environment that enables promoter-proximal pausing, and opens the possibility for a novel role of BEAF-32 and Chromator as pausing factors.

**Figure 4:**
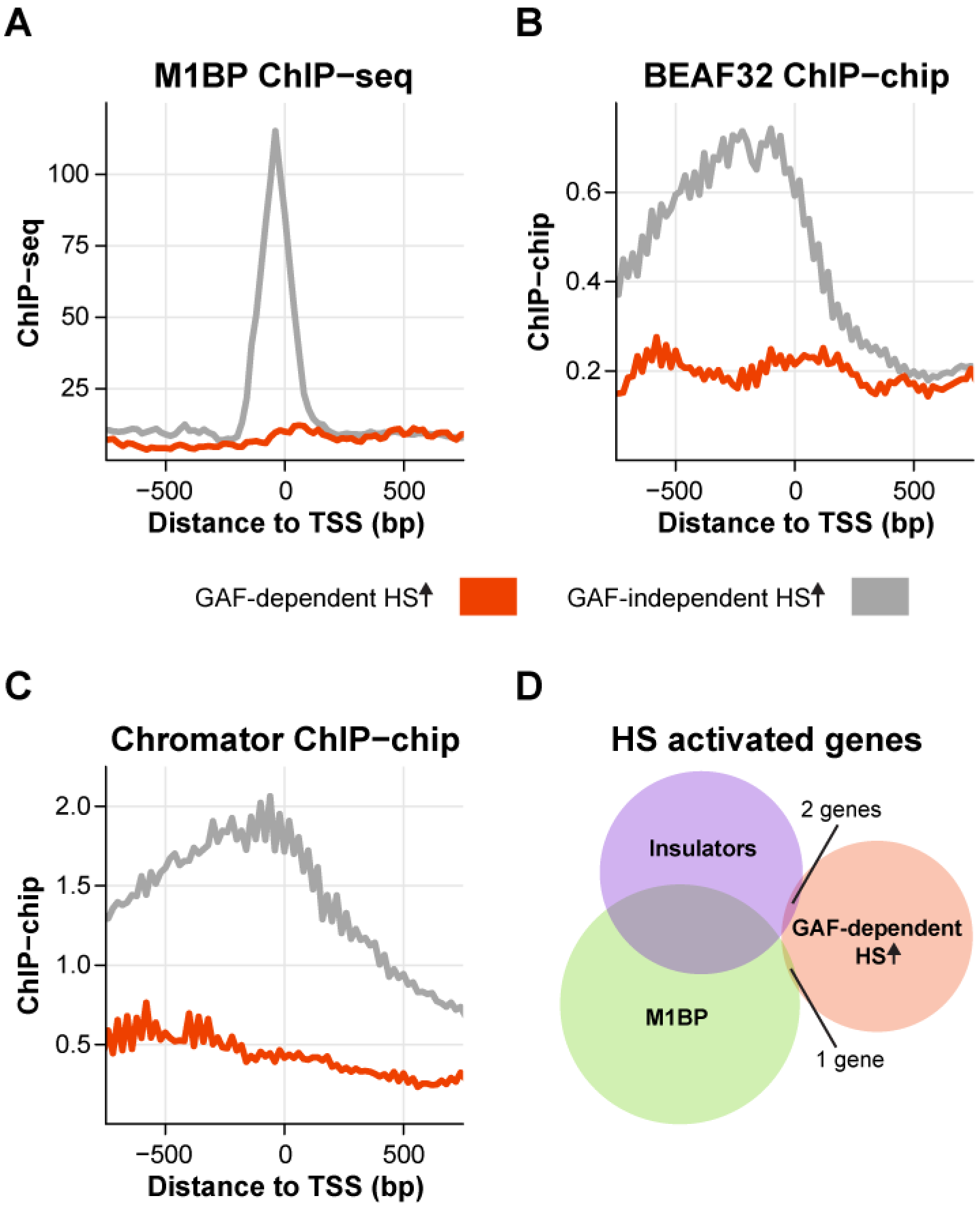
Insulator proteins and M1BP are enriched in the promoter region of HS activated genes with GAF-independent induction. **(A-C)** Average M1BP ChIP-seq **(A)**, BEAF-32 ChIP-chip **(B)**, and Chromator ChIP-chip **(C)** signal between −750 to +750 bp to the TSS (in 20 bp bins) of genes with GAF-dependent (n=44) or GAF-independent (n=199) HS activation. **(D)** Venn diagram showing the overlap between HS activated genes with GAF-dependent activation and genes bound by M1BP or both insulator proteins (BEAF-32 and Chromator) within ±1kb of the TSS.

### HSF is essential for the induction of only a small minority of HS activated genes

HSF is the evolutionarily conserved master regulator of the HS response and is essential for the activation of classical HSP genes (Wu 1995). Inducible HSF binding at those genes is critical for the recruitment of the positive elongation factor P-TEFb (Lis et al. 2000), which modulates the release of Pol II into productive elongation. We used our previously published HSF ChlP-seq datasets, performed before and after 20min of HS induction (Guertin and Lis 2010), to determine if HSF also preferentially binds to non-canonical HS activated genes. HSF ChIP-seq peaks are closer to the TSS in the HS activated class when compared to the repressed (Figure 5A, Kolmogorov-Smirnov test p-value = 4.04 × 10^-12^) and unchanged gene classes (Figure 5A, Kolmogorov-Smirnov test p-value = 1.6 × 10^-7^). Surprisingly, even though HSF is enriched in the proximity of activated genes, less than 20% of those genes have an HSF ChIP-seq peak within ±1kb of the TSS. The existence of HSF-independent genes has been previously demonstrated in *Drosophila* (Gonsalves et al. 2011). However, our study substantially expands the number of identified genes and offers a more comprehensive view of HSF-independent regulation due to the considerably higher resolution and sensitivity afforded by our binding and nascent transcription assays. These results indicate that HSF can activate genes when bound to distal enhancer sites or that there are other factors dictating the induction of HS activated genes.

**Figure 5:**
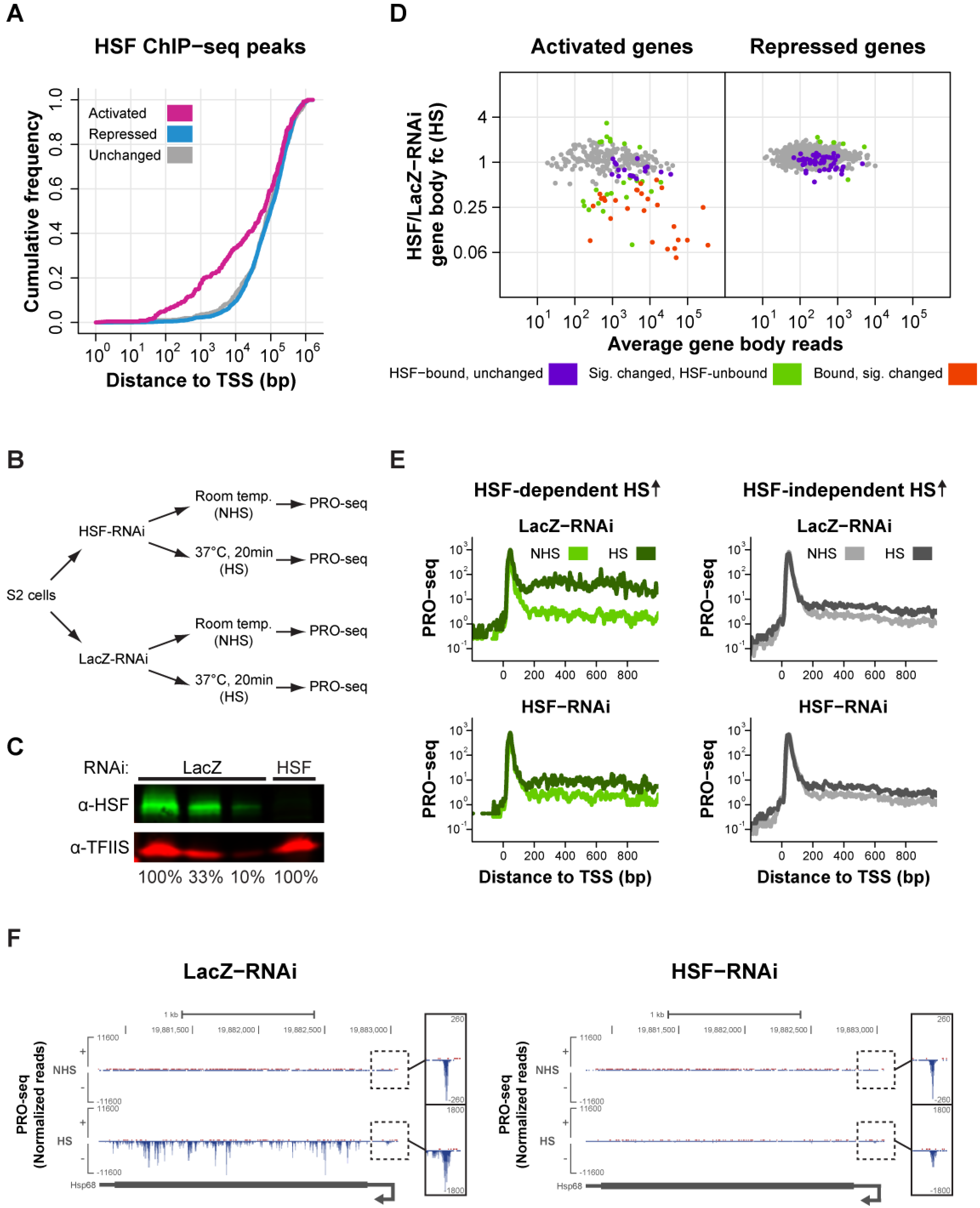
HSF is essential for the induction of only a small minority of HS activated genes and activates genes by stimulating the release of paused Pol II. **(A)** Cumulative distribution plots of the distance between the closest HSF ChIP-seq peak and the TSS of each gene in the HS activated (n=249), repressed (n=2300), and unchanged (n=517) classes. **(B)** Experimental set-up. *Drosophila* S2 cells were treated with either HSF or LacZ RNAi for 5 days. Nuclei were then isolated for PRO-seq after cells were incubated at room temperature (NHS) or <u>H</u>eat <u>S</u>hocked for 20 minutes (HS). **(C)** Western blot of whole cell extracts from LacZ and HSF-RNAi treated cells for HSF and TFIIS (loading control). 100% is equivalent to 1.5 × 10^6^ cells. **(D)** DESeq2 analysis to determine the effect of HSF-RNAi treatment on the PRO-seq gene body reads after HS for the HS activated and repressed classes. We used DESeq2 to identify significantly changed genes between HSF-RNAi HS and LacZ-RNAi HS cells and the results are displayed as MA plots. Significantly changed genes were defined using an FDR of 0.001. HSF-bound genes are labeled in purple, significantly changed genes (according to DESeq2) are labeled in green and genes that are both HSF-bound and significantly changed are labeled in orange. fc = fold-change. **(E)** Average normalized PRO-seq reads between −200 to +1000 bp to the TSS (in 5 bp bins) of genes with HSF-dependent (n=44) or independent (n=197) HS activation for the LacZ-RNAi and HSF-RNAi datasets before (NHS) and after HS. **(F)** Representative view in the UCSC genome browser (Kent et al. 2002) of a gene with HSF-dependent activation. PRO-seq normalized reads for the different RNAi treatments (LacZ and HSF) before and after HS for the plus strand are shown in red and for the minus strand in blue. Gene annotations are shown at the bottom.

### HSF activates genes by stimulating the release of paused Pol II

To investigate HSF's roles during the HS-induced transcriptional response, we performed PRO-seq in biological replicates of HSF-RNAi treated cells prior to HS and after 20 minutes of HS (Figure 5B–C, Table S1) (Spearman's coefficient ranged between 0.96-0.99, Figure S1A-B, middle panels). Comparison of the HS gene body reads in the LacZ-RNAi control and HSF-RNAi for activated and repressed genes shows that the HS PRO-seq levels of 20% of activated genes were affected by HSF-RNAi, while a significant change was only observed for <1% of the repressed class, demonstrating that HSF is important for HS activation, but not for repression (Figure 5D).

Most of the activated genes that have HSF binding within ±1kb of the TSS have reduced HS induction after HSF knockdown (Figure 5D, left panel, orange points). HSF-bound genes with compromised induction upon HSF knockdown are enriched for HSF binding immediately upstream (within 200 bp) of the TSS, while the unaffected class displays a random distribution of distances (Figure S6A). Furthermore, HSF-bound genes with reduced HS induction have significantly higher HSF ChIP-seq binding intensity when compared to the unaffected class (Figure S6B, Mann-Whitney *U* test p-value = 2.5 × 10^−3^), indicating that higher HSF binding levels and positioning upstream and proximal to the TSS are important for the induction of HSF's target genes. Additionally, comparison of all induced, HSF-dependent genes to the remainder of HS-activated genes (HSF-independent HS activation) showed that genes depending on HSF have stronger HS induction (Figure 5E).

HSF depletion does not affect the induction of most genes that are not bound by HSF within 1kb of the TSS (Figure 5D, left panel, gray points). Nonetheless, the presence of significantly changed genes with no proximal HSF binding (Figure 5D, left panel, green points) indicates that HSF may be able to mediate activation at distal enhancer sites on a small subset of genes. The enhancer activity of HSF had been previously shown in a focused study of *Hsp70* to be weak and require large arrays of HSF binding sites (Bienz and Pelham 1986). This early study and the rarity with which we find HSF acting at a distance might be explained if such long-range interactions required specialized binding sites and chromatin architecture. Clearly, the preferred mode of HSF action is close to promoters.

Composite profiles show that the average pausing levels of genes with HSF-dependent and HSF-independent HS activation are not affected by HS in both LacZ-RNAi and HSF-RNAi conditions (Figure 5E), indicating that neither HSF depletion nor HS have much of an effect on pausing. Quantification of the effect of HSF knockdown on pausing levels in both NHS and HS conditions for all HS activated genes revealed that the pausing level of only one gene was significantly affected by the knockdown (Figure S6C). Figure 5F has an example of a HS-activated gene in which HSF knockdown does not have a significant effect on pausing levels but substantially inhibits HS induction. Taken together, these results suggest that HSF acts mainly at the release of paused Pol II into productive elongation, which is consistent with the critical role of HSF in the recruitment of the pause release factor, P-TEFb, to *Hsp70* (Lis et al. 2000).

### HS transcriptional repression is HSF-independent and results in a decrease of promoter-proximally paused Pol II

Our data revealed that HS induction causes a vast transcriptional shutdown, with thousands of genes being repressed after 20min of increased temperatures (Figure 1A). To elucidate the mechanisms involved in this repression, we observed the PRO-seq profile for all repressed genes plotted as heatmaps before and after HS (Figure 6A). This analysis indicates the presence of enriched PRO-seq reads in the region immediately downstream of the TSS, representing promoter-proximally paused polymerases. Both gene body reads and reads in this promoter-proximal region are reduced after HS, which is also evidenced by the overall blue color of the fold-change heatmap (Figure 6A, right panel). The overall NHS and HS distributions and the HS/NHS fold-change are very similar after HSF depletion by knockdown (Figure 6B), indicating that HSF does not play a role in gene repression by HS.

**Figure 6:**
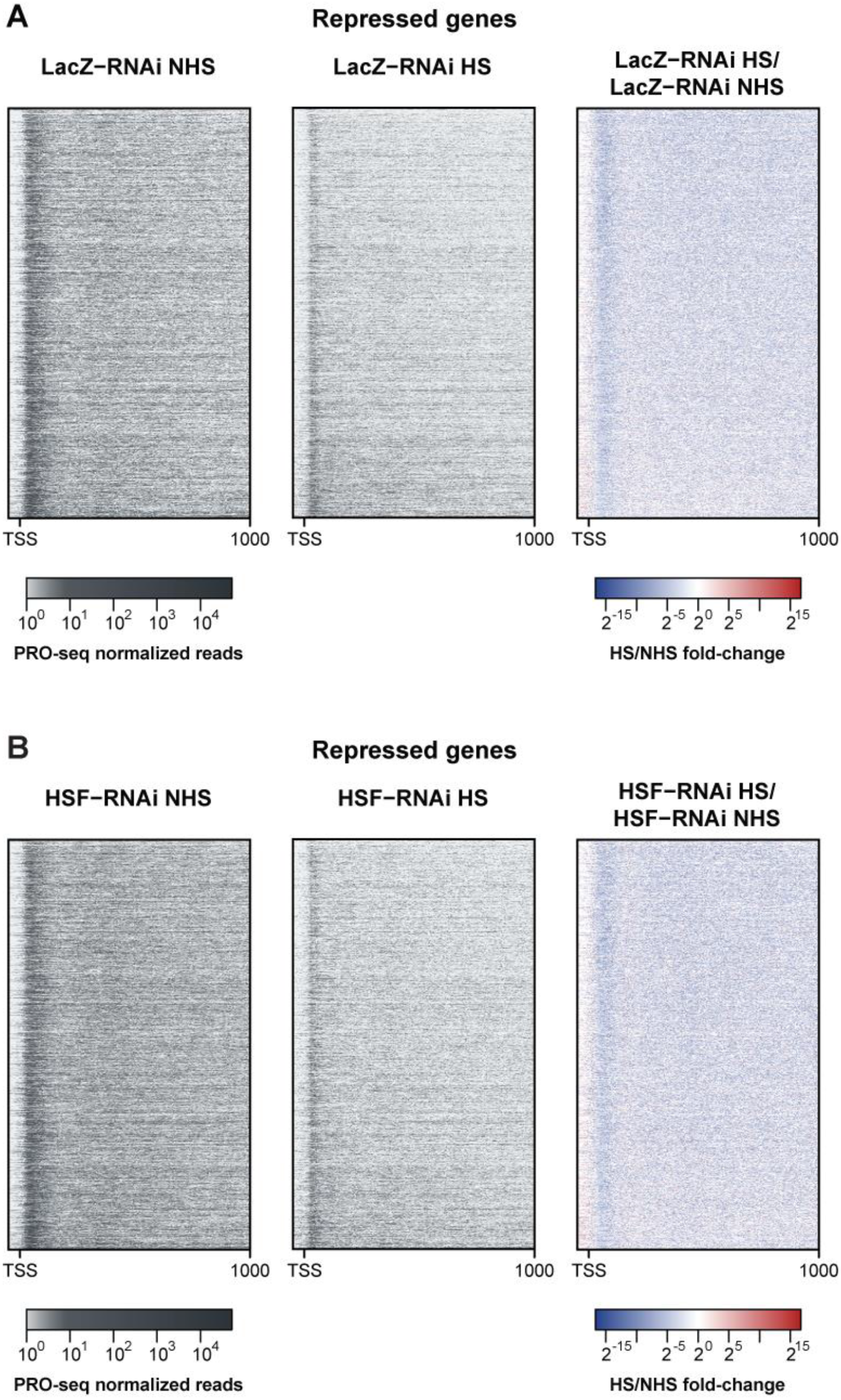
HS transcriptional repression is HSF-independent and results in a decrease of promoter-proximally paused Pol II. Heatmaps showing the NHS PRO-seq levels (left panel), HS PRO-seq levels (middle panel) and the fold-change between the two conditions (right panel) between −1000 to +1000 bp to the TSS (in 5 bp bins) of HS repressed genes (n=2300) for the LacZ-RNAi **(A)** and HSF-RNAi **(B)** treatments. Genes in both A and B are sorted by the HS/NHS PRO-seq fold-change in the LacZ-RNAi condition (highest to lowest).

## Discussion

In this study, we used PRO-seq to comprehensively characterize the direct changes in Pol II distribution that occur in *Drosophila* S2 cells in the minutes following HS. We show that the HS response is more general than previously appreciated, with thousands of genes being repressed and hundreds activated by heat stimulus. This latter class is not limited to the group of cellular chaperones that are known to be activated by stress (Lindquist and Craig 1988), and includes hundreds of other genes with various cellular functions (Table S2). Surprisingly, only a minority of the activated genes are regulated by HSF, which was previously believed to be the major orchestrator of the response. Moreover, we show that promoter-proximal pausing is highly pronounced and prevalent among activated genes prior to HS. GAF, which has been shown to be important for establishing pausing, is highly enriched at the promoter of HS activated genes, and our results suggest that GAF-mediated pausing in a subset of these genes is essential for HS activation. Furthermore, our results indicate that HS repression is regulated at the step of transcription initiation in *Drosophila*, and this process is independent of HSF. Overall, by measuring how transcription changes after HS, our results provide insights into mechanisms of transcription activation and repression, the key regulating factors and the steps in the transcription cycle that are modulated.

### GAF-mediated promoter-proximal pausing is essential for the HS activation of a subset of genes

Classical HSP genes accumulate paused Pol II molecules between 20-50 bp downstream of the TSS prior to HS (Rougvie and Lis 1988; Rasmussen and Lis 1993). Our results and analyses greatly expand upon these previous findings and indicate that pausing is a common feature among HS activated genes and is not specific to the highly induced class of molecular chaperones. Previous studies have shown that the paused Pol II complex on *Hsp70* and many other genes is remarkably stable (Henriques et al. 2013; Buckley et al. 2014; Jonkers et al. 2014), and this stably paused molecule can help to maintain an open chromatin environment that is accessible to transcription factors that will promote the release of Pol II into productive elongation, mediating a rapid response to HS. The open chromatin state of our newly identified HS activated genes is confirmed with the higher DNase I hypersensitivity signal observed in the promoter region relative to repressed and unchanged genes (Figure S4A).

GAF has been previously shown to be important for establishing pausing and is highly enriched in the promoter region of activated genes prior to HS. GAF is essential for HS activation of a subset of GAF-bound genes that have high levels of GAF binding in the region immediately upstream of the core promoter, indicating that GAF's positioning and levels are important for its role in the HS response. We also observed that the pausing levels of genes with GAF-dependent HS activation are dramatically reduced upon GAF depletion prior to HS. Transgenic studies of the model *Hsp70* gene have demonstrated that presence of the GAF binding element is essential for generating pausing at this gene and that pausing level changes created by mutating the core promoter strongly correlate with the promoter's potential to induce transcription upon HS induction (Lee et al. 1992). Our results expand upon these studies and demonstrate that GAF depletion prior to HS in the native chromatin environment of a subset of HS activated genes abrogates Pol II pausing levels and the consequent induction of these genes by HS. Importantly, other GAF-bound genes that maintain pausing upon GAF knockdown, presumably due to the activity of other pausing factors like M1BP and the insulator protein BEAF-32, remain fully HS inducible. We propose a model where GAF-mediated pausing is essential to maintain an open chromatin environment at the promoter region prior to HS (Figure 7A). When GAF is depleted by knockdown and pausing is not properly established, then the promoter loses its potential to induce transcription after HS (Figure 7A).

**Figure 7:**
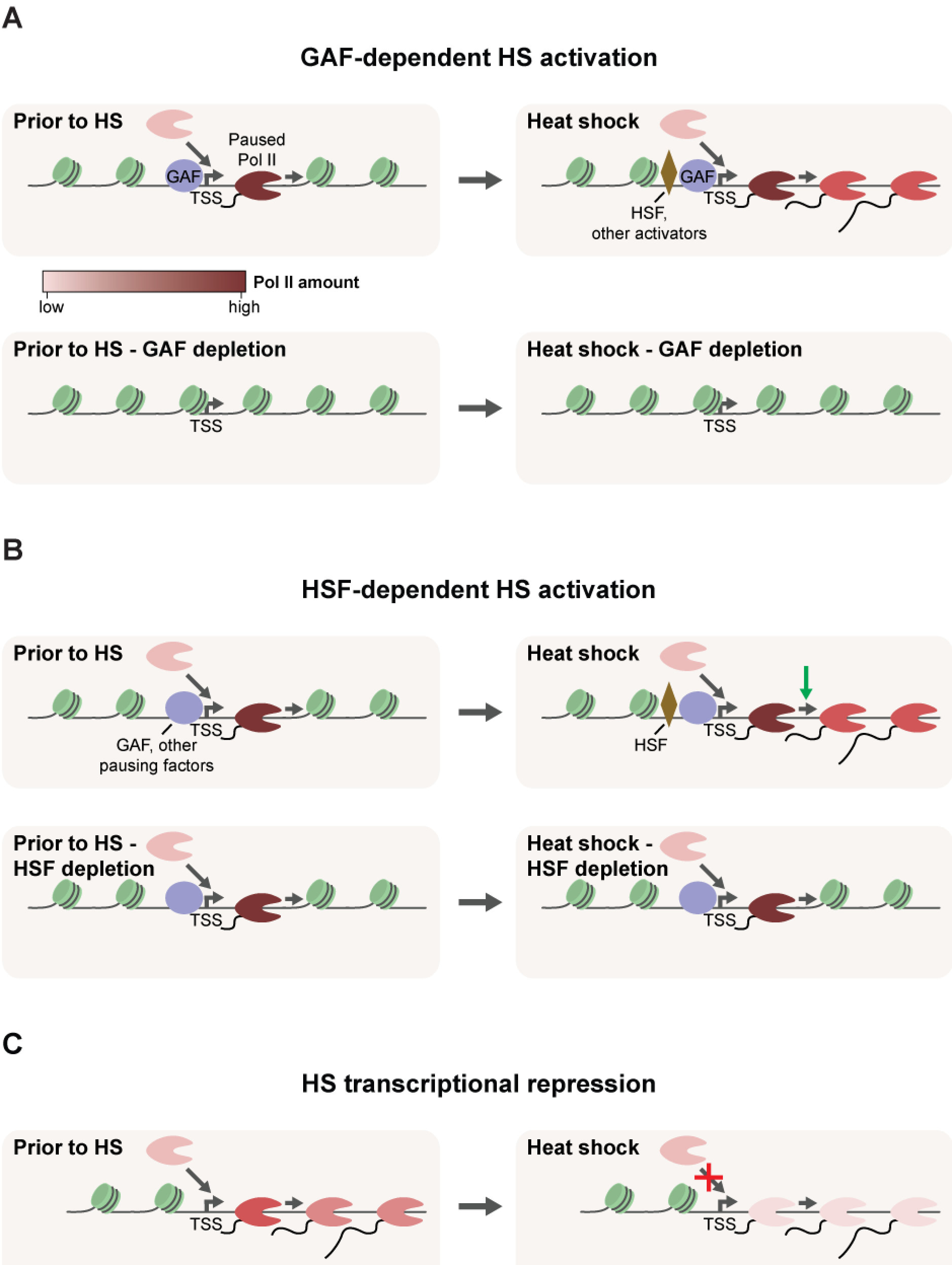
Summary of proposed mechanisms of HS transcriptional regulation. Model depicting the mechanisms of transcriptional regulation proposed in our study for **(A)** GAF-dependent HS activation, **(B)** HSF-dependent HS activation and **(C)** HS transcriptional repression. Red X represents a step that is being inhibited, and green arrow represents a step induced by HS. Nucleosomes are shown in green.

### HSF acts at the step of promoter-proximal pausing release

HSF depletion has almost no effect on pausing reads both before and after HS (Figure S6C), and the average pausing levels are largely unaffected by HS and HSF knockdown (Figure 5E). The amount of pausing is determined by the transcription recruitment/initiation rate and the rate of escape into productive elongation. If HSF was acting at the step of Pol II initiation, we would expect the pausing levels to be reduced upon HSF depletion, which is not observed. Therefore, we propose a model where after being recruited to the promoter region upon HS, HSF promotes the release of Pol II into the gene body, likely through the indirect recruitment of P-TEFb, which has been shown to be the case for classical HSP genes (Lis et al. 2000). Pausing also maintains an open chromatin environment that is accessible to transcription activators such as HSF. In this model, the activity of factors that are important for establishing pausing prior to HS, such as GAF and possibly M1BP and BEAF-32, is crucial for HSF-dependent HS activation, and failure to generate pausing prevents the induction of HSF target genes.

### HS causes a rapid and broad reduction in transcription, which is regulated at the transcription initiation step and independent of HSF

Early low resolution studies in *Drosophila* polytene chromosomes have shown that HS causes a genome wide downregulation of transcription (Spradling et al. 1975; Jamrich et al. 1977), presumably to reduce the accumulation of misfolded protein aggregates. Although this has been a paradigm in the HS field, higher resolution genome-wide studies have failed to identify all the primary genes that are repressed by heat, mostly due to the limitations of measuring steady-state levels of mRNA, which requires that the mRNAs already present in the cells have shorter half-lives than the HS time points used in the experiment. Our results provide definitive evidence to support the widespread shutdown of transcription caused by HS. We identify and quantify the genes with significantly reduced transcription and demonstrate that the HS repressive response is very rapid, with over a thousand genes being repressed after only 5 minutes of HS (Figure S3A). Furthermore, the Pol II density in the promoter-proximal region, which represents the paused Poll molecules, is also significantly reduced across all HS repressed genes (Figure 6). The accumulation of Pol II in the pausing region depends on both the transcription initiation rate and the rate of escape into productive elongation. The reduction in pausing levels thus indicates that the recruitment and initiation of Pol II is affected by HS (Figure 7).

The HS-induced binding of HSF is not essential for the genome-wide transcriptional repression (Figure 6), and given the magnitude of this repressive response, we believe that it is unlikely that one single transcription repressor is responsible for inhibiting transcription initiation in all HS repressed genes. We believe that there are three major possible mechanisms that could be responsible for HS-mediated repression. (1) The activity of a general transcription factor that is involved in recruitment of Pol II to the promoter could be modulated by heat stimulus. (2) Changes in nucleosomal composition or positioning induced by heat could generate an unfavorable chromatin environment that would prevent transcription initiation and elongation. Indeed, a previous study has demonstrated that HS results in decreased nucleosome turnover genome-wide within gene bodies (Teves and Henikoff 2011). However, the same pattern was observed after drug inhibition of Pol II elongation, which argues that reduced nucleosome turnover may be a consequence rather than the cause of the genome wide transcriptional repression (Teves and Henikoff 2011). (3) A genome-wide rearrangement of the 3D chromatin structure could mediate transcriptional repression, either by disrupting long-range interactions that are needed for transcription prior to HS or by allowing new long-range interactions that repress transcription initiation. In support of the latter possibility, a recent study in a different *Drosophila* cell line demonstrated that HS induces a genome-wide rearrangement in the 3D nuclear architecture through relocalization of architectural proteins from <u>T</u>opologically <u>A</u>ssociated <u>D</u>omain (TAD) borders to inside TADs, which results in an increase in long-distance inter-TAD interactions (Li et al. 2015). The promoter contacts generated by these newly formed long-range interactions are enriched for Polycomb complexes, arguing for a role of Polycomb in HS transcription repression (Li et al. 2015). This model suggests inhibition of the transcription initiation that we observe could be mediated by new Polycomb contacts. These three possible mechanisms are not mutually exclusive. However, any model must accommodate our new observations that 1) recruitment of Pol II is the step in the transcription cycle that is regulated, 2) HSF is not involved in the repression; 3) the specifically repressed genes identified here and their level of down-regulation must be accommodated by any proposed regulatory factor interactions.

## Materials and methods

### GAF, HSF and LacZ RNAi treatments

*Drosophila* S2 cells were grown in M3+BPYE media supplemented with 10% FBS until they reached 3-5 × 10^6^ cells/mL. At this point, the cells were split to 1 × 10^6^ cells/mL in serum-free M3+BPYE media, and the desired volume of cells was mixed with LacZ, HSF or GAF dsRNA to a final concentration of 10 μg/mL. After incubation at 25°C for 45 minutes, an equal volume of M3+BPYE media supplemented with 20% FBS was added to the cells. After 2.5 days, the cells were split 1:2 into two new flasks and more dsRNA was added to keep the final concentration at 10 μg/mL. After 2.5 days the cells were HS treated and harvested for nuclei isolation.

The dsRNAs used in these experiments were transcribed from a dsDNA template that had a T7 polymerase promoter at both ends. The DNA templates were generated by PCR using the following primers:

LacZ Forward: GAATT AAT ACGACTCACTATAGGGAGAGATATCCTGCTGATGAAGC
LacZ Reverse: GAATT AAT ACGACTCACTATAGGGAGAGCAGGAGCTCGTTATCGC
GAF Forward: GAATTAATACGACTCACTATAGGGATGGTTATGTTGGCTGGCGTCAA
GAF Reverse: GAATT AAT ACGACTCACTATAGGGATCTTT ACG CGTGGTTT GCGT
HSF Forward: GAATTAATACGACTCACTATAGGGAGAGCCTTCCAGGAGAATGCA
HSF Reverse: GAATTAATACGACTCACTATAGGGAGAGCTCGTGGATAACCGGTC

### Western blot analysis to assess knockdown levels

Western blots were performed using standard conditions, and dilutions of the LacZ-RNAi control sample were used as a quantitative indication of signal linearity. Lab stocks of rabbit anti-HSF and anti-GAF antibodies and guinea pig anti-TFIIS antibody were used at dilutions of 1:2000, 1:500, and 1:3000, respectively. We used IRDye 800CW donkey anti-rabbit (1 mg/mL) and IRDye 680LT donkey anti-guinea pig (1 mg/mL) as secondary antibodies at a 1:15000 dilution and the membrane was imaged using the LI-COR Odyssey imaging system.

### Heat Shock treatments

For the HS treatments, an equal volume of M3+BPYE medium (no serum) at 48°C was added to the cells, and the cultures were incubated at 37°C for the desired time.

### Preparation of PRO-seq libraries

Nuclei isolation and PRO-seq library preparation were performed as described previously (Kwak et al. 2013).

### Preparation of RNA-seq libraries

Total RNA from S2 cells was extracted using TRIzol reagent (Thermo Fisher Scientific) and then isolated from the aqueous phase using the E.Z.N.A. Total RNA Kit I (Omega Bio-tek). The following steps were performed by the Cornell RNA Sequencing Core (Department of Biomedical Sciences, College of Veterinary Medicine, Cornell University). PolyA+ RNA was isolated with the NEBNext Poly(A) mRNA Magnetic Isolation Module (New England Biolabs). TruSeq-barcoded RNA-seq libraries were generated with the NEBNext Ultra Directional RNA Library Prep Kit (New England Biolabs). Each library was quantified with a Qubit 2.0 (dsDNA HS kit; Thermo Fisher Scientific) and the size distribution was determined with a Fragment Analyzer (Advanced Analytical) prior to pooling.

### PRO-seq data acquisition

PRO-seq libraries were sequenced in 50 nt runs on the Illumina HiSeq, using standard protocol at the Cornell Biotechnology Resource Center (http://www.BRC.cornell.edu). Raw sequencing reads were processed using the FASTX-Toolkit (http://hannonlab.cshl.edu/fastx_toolkit/index.html). Illumina adapters were removed with the fastx_clipper tool and reads were trimmed to 26-mers using fastx_trimmer. Sequencing reads shorter than 15 nucleotides were discarded. fastx_reverse_complement was then used to generate the reverse complement of the sequencing reads, which correspond to the sense strand of nascent RNA in the nucleus. Reads were aligned uniquely to the *Drosophila melanogaster* dm3 reference genome using Bowtie (Langmead et al. 2009) with up to two mismatches. Histograms of the 3′-end position of each mapped read in base-pair resolution were generated in bedgraph format and used for all subsequent analyses. Table S1 contains a summary of sequencing yields and the number of reads that mapped uniquely to the genome or other annotations. Replicates were highly correlated and were pooled for further analyses (Figure S1). Sequencing datasets can be found under GEO accession number GSE77607.

### PRO-seq normalization method

We used a previously published Pol II ChIP-seq dataset in *Drosophila* S2 cells (Teves and Henikoff 2011) to identify genes whose transcription does not change during HS. Unlike ChIP-seq reads, which can originate from both sense and anti-sense strands, PRO-seq reads are strand specific. Therefore, in order to increase the likelihood of selecting genes that have the majority of their reads originating from the sense strand, we used our PRO-seq LacZ-RNAi control datasets (NHS and 20min HS) to identify and filter out genes that have high levels of transcription in the anti-sense strand. To identify those genes, we calculated the fraction of PRO-seq reads originated from the anti-sense strand for each gene and only kept the ones whose fraction is less or equal than 0.2 for both the NHS and 20min HS conditions. Because of the high background in ChIP-seq data, we then focused on genes with highest levels of ser2-P ChIP signal (*Z* score > 3) (Core et al. 2012), assuming these will contain the highest densities of transcribing Pol II over background. In order to obtain a final subset of unaffected genes, we filtered out the ones whose gene body fold-change between NHS and HS conditions is less than 0.85 and greater than 1.15, resulting in 335 genes. The mRNA levels of this subset of genes are also unaffected after HS (Figure S7), which is consistent with the transcription levels of these genes not changing after induction.

We then used the sum of the total number of gene body reads for all 335 genes to generate normalization factors in our PRO-seq data to normalize the datasets between replicates and different time points. Since the GAF and HSF RNAi treatments did not result in genome-wide changes in transcription in both the NHS and 20min HS time points, we used the same subset of 335 unaffected genes to normalize the datasets between different RNAi treatments (LacZ, GAF and HSF).

To access the efficacy of this normalization method, we examined the correlation between gene body and promoter reads for replicates (Figure S1) and gene body reads across different time points (Figure S8A-B). All replicates show good correlations and time points that are closer to each other have higher correlation coefficients and better fits to the 1:1 diagonal. Moreover, for the RNAi treatments, we examined the correlation between gene body reads across different conditions (NHS and 20min HS) and observed that the NHS datasets have higher correlation coefficients and better fits to the 1:1 diagonal when plotted against each other, and the same was observed for the HS treatments (Figure S8C-D). Taken together, these results indicate that the normalization method worked appropriately.

### Differential expression analysis using DESeq2

We used DESeq2 (Love et al. 2014) to identify genes whose gene body reads significantly change after HS, starting with a list of 9452 non-overlapping genes described previously (Core et al. 2012). Gene body reads were collected from 200 bp downstream of the TSS, and we used different 3′ limits for each time point, assuming a conservative estimate for Pol II transcription elongation rate of 1kb/min. We provided our own normalization factors for the DESeq2 calculations, which were obtained as described above. We used an FDR of 0.001 to identify activated and repressed genes. Unchanged genes were defined as having an adjusted p-value higher than 0.5 and log_2_fold-change higher than —0.25 and lower than 0.25.

### Upstream transcription filter

Due to the compactness of the *Drosophila* genome, some of the genes identified as differentially expressed by DESeq2 appear to be false positives caused by changes in run-through transcription originated at the upstream gene. To minimize the number of false positives, we implemented a filter to exclude from our analyses genes that have high levels of transcription in the region immediately upstream of the TSS. For each gene, we obtained the read counts from a window upstream of the TSS (−500bp to −100bp of the TSS) and a window in the gene body (+300bp to +700bp of the TSS) (Figure S9A). The 3′ limit of the upstream window was defined as −100bp to the TSS to avoid confounding effects of potentially misannotated TSSs. In the case of the gene body window, the 5′ limit was defined as +300bp to the TSS to avoid the region immediately downstream of the TSS, which can contain peaks of promoter-proximal paused Pol II. We then took the ratio of the read counts in these two regions for each gene, taking into consideration the number of mappable positions in the two windows (Figure S9A). This ratio was named *upstream ratio* and was later used to exclude false positive genes from our analyses.

In order to verify if we could distinguish true and false positives based on the upstream ratio and define the appropriate cutoff to filter out false positive genes, we visually inspected 100 randomly selected activated genes and classified each one as true or false positive based on the presence or absence of run-through transcription. The average mRNA levels (Figure S9B) are higher for the genes that were defined as true positives, which provides an independent verification of the criteria that were used to define true and false positives.

The distribution of upstream ratios for the LacZ-RNAi HS condition was very distinct for true and false positives, with very little overlap (Figure S9C), indicating that this metric could be used to identify false positives. In order to define the optimal cutoff, we evaluated the performance of all potential cutoffs from 0 to 1 in 0.01 increments and used the accuracy metric ((true positives + true negatives)/total) to identify the cutoff with the best performance (Figure S9D). The horizontal line in Figure S9C represents the chosen cutoff (0.23). Filtering out genes with upstream ratios greater than 0.23 eliminates all but one false positive, with only a minor loss of true positive genes.

We then filtered out genes with upstream ratio higher than 0.23 in the activated and repressed classes to generate the final subsets of genes that were used in all subsequent analyses. In order to use the condition with highest PRO-seq levels for upstream ratio calculation, we used the NHS upstream ratio to filter the repressed subsets of genes and the HS upstream ratio to filter the activated subsets for every HS and RNAi treatment.

### Gene ontology analysis

Gene ontology analysis on HS activated and repressed genes were performed using the Functional Annotation tool from DAVID (Huang et al. 2009b, 2009a), in which ‘GOTERM_BP_FAT’ was selected.

### Promoter-proximal pausing analysis

The “pausing region” was defined as the 50bp interval with highest number of reads within −50 to +150bp of the TSS. This region was defined using the LacZ-RNAi control NHS condition and the same interval was used for all the other treatments and conditions. Pausing index was then calculated as the ratio of the read density in the pausing region (reads/mappable bases) and the read density in the gene body (as defined above). Genes were classified as paused as described previously (Core et al. 2008). We used DESeq2 to identify genes whose pausing levels significantly change after HS, using an FDR of 0.001.

### Distance analysis

We used *bedtools closest* (Quinlan and Hall 2010) to identify the closest HSF, GAF or M1BP ChIP-seq peak to the TSS of every transcription unit in our list. HSF, GAF or M1BP-bound genes were defined as having a ChIP-seq peak within ±1000 bp of the TSS. The GAGA element was identified de novo in the promoter region of activated genes (−300 to +50 bp of the TSS) using *MEME* (Bailey and Elkan 1994).

### RNA-seq data acquisition and analysis

RNA-seq libraries were sequenced in 100 nt runs on the Illumina HiSeq, using standard protocol at the Cornell Biotechnology Resource Center (http://www.BRC.cornell.edu). Illumina adapters were removed with the fastx_clipper tool (http://hannonlab.cshl.edu/fastx_toolkit/index.html) and sequencing reads shorter than 20 nucleotides were discarded. Reads were aligned to the *Drosophila melanogaster* dm3 reference genome/transcriptome using TopHat2 (Kim et al. 2013), with the following parameters: “--library-type fr-firststrand --no-novel-juncs” Table S3 contains a summary of sequencing yields and the number of reads that mapped to the genome/transcriptome or other annotations. Sequencing datasets can be found under GEO accession number GSE77607.

FPKM values for each gene were generated with Cuffnorm (Trapnell et al. 2010), using the BAM files generated by TopHat2 as input. Raw counts for each gene were obtained using HTSeq-count (Anders et al. 2014) and used as input for differential expression analysis using DESeq2 (Love et al. 2014). We used an FDR of 0.001 to identify genes whose mRNA levels significantly increase or decrease upon HS.

## Acknowledgements

We thank Jian Li and David Gilmour for providing the M1BP ChIP-seq peak information file. We also thank current and past members of the Lis lab for insightful discussions and comments on the manuscript.

## Supplemental Material

**Figure S1:** Biological replicates of PRO-seq libraries were highly correlated for both promoter and gene body regions.

**Figure S2:** Measurement of steady-state mRNA levels by RNA-seq is unable to detect a genome-wide shutdown of transcription after HS.

**Figure S3:** Substantial genome-wide response to HS occurs as early as 5 minutes post-HS.

**Figure S4:** Promoter region of HS activated genes is more accessible than repressed and unchanged classes prior to HS.

**Figure S5:** Higher binding levels and positioning immediately upstream of the core promoter are important for GAF's role in HS activation.

**Figure S6:** Higher HSF binding levels and positioning upstream and proximal to the TSS are important for the induction of HSF's target genes.

**Figure S7:** mRNA levels of 335 genes used for normalization are not affected by HS.

**Figure S8:** Validation of the PRO-seq normalization method used in our study.

**Figure S9:** Validation of the upstream transcription filter implemented in our study.

**Table S1:** Sequencing and alignment of PRO-seq libraries.

**Table S2:** Identification and quantification of genes that are significantly activated or repressed after 20min of HS.

**Table S3:** Sequencing and alignment of RNA-seq libraries.

**Table S4:** Pausing index calculation and identification of paused genes.

